# SLAST: Simple Local Alignment Search Tool

**DOI:** 10.1101/840546

**Authors:** Juanjo Bermúdez

## Abstract

We present a local alignment search tool not based on the usual strategy of seed and grow often employed for these tools. Instead, we just find regions in the database sequences having a high density of seed matches and then we perform a Smith-Waterman local alignment of the query sequence into these regions. This approach has some advantages for some use cases.

## I. INTRODUCTION

BLAST [1] is by far the most widely used tool for rapid similarity searching among nucleic acid or amino acid sequences.

BLAST does well its job and there are also many alternatives that apparently work fairly well. Therefore, there is no practical reason to justify why we developed SLAST. We just had the opportunity to develop it and we did. Indeed, to be honest, we developed it because we were unable to make BLAST work properly due to bad parameter selection. We found later that it works well with the right parameters but we already had developed SLAST by that time.

Therefore, the basic idea of SLAST does not have, in practice, any big advantage over BLAST or other similar tools. But we think the way in which it is encapsulated to provide a functionality does indeed have some advantages, and despite the core technology of SLAST is in theory replaceable by any of the other ones (which in theory are faster and with similar reliability), not doing that does not impose any big impediment for the functionality of the tool. We are not sure either that there are theoretical reasons why SLAST could not evolve to be as fast as BLAST. It could be that it actually isn’t because of a lack of optimization as it is just a first version.

What SLAST actually really does differently than BLAST (not sure about the other ones) is the formatting of the results. SLAST is really(!!) a local alignment search tool, which BLAST is not. SLAST makes a search for the complete query sequence, not for subsequences of the query sequence. The output of SLAST are really local alignments of the full query sequence into the database sequences. Indeed, the output are alignments done by means of the Smith-Waterman [2] algorithm.

We observe that finding local alignments instead of similar subsequences has some practical advantages that are actually not easily available for BLAST users.

## II. METHOD

As has already been explained, the idea behind SLAST is quite simple: finding segments of sequences in the database with a high density of seed matches and then applying Smith-Waterman to these regions. These are the steps involved:

### 1. Selecting seeds

There are some possible configurations. For example, we can split the query sequence into same-size seeds or we can select as seeds every subsequence of the query sequence. We can also implement intermediate strategies, like selecting subsequences of fixed length starting at even positions of the query sequence. Depending on this configuration the algorithm will go faster or slower and will have more or less accuracy.

The way in which the database is indexed also affects the performance of the algorithm. If we index the sequences splitting them into same-size chunks starting at even positions it will, in theory, miss some matches, but using every position as start it will cover all possibilities (at the cost of more space needed for indexing).

**Fig. 1.**
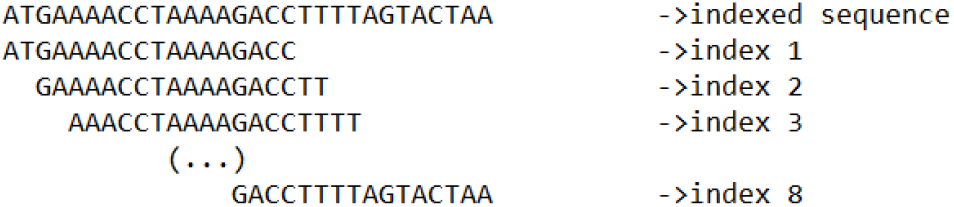
Indexing based on same-size chunks (16 bps) located at even positions.

**Fig. 2.**
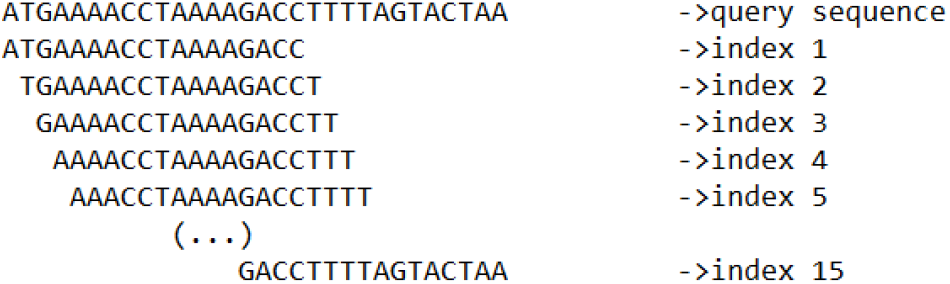
Indexing based on same-size chunks (16 bps) located at every position.

### 2. Making a search for every seed

The indexing technology that we use is able to match sequences even if they have some characters deleted, inserted, or modified. Therefore, even if the indexing configuration does not cover all subsequences in the database, it could anyway find the corresponding matches. The sensibility to mismatches, insertions or deletions can also be configured in another parameter. The user only needs to keep in mind that if he uses that ability in the indexing technology to cover possible misses, then it will miss some matches corresponding to insertions or deletions that really occurred by evolution (as the ‘wildcard’ will be used to cover the artificial miss or misses generated by the configuration). You also have to think that the more sensible the indexing technology is, the slower it is.

**Fig. 3.**
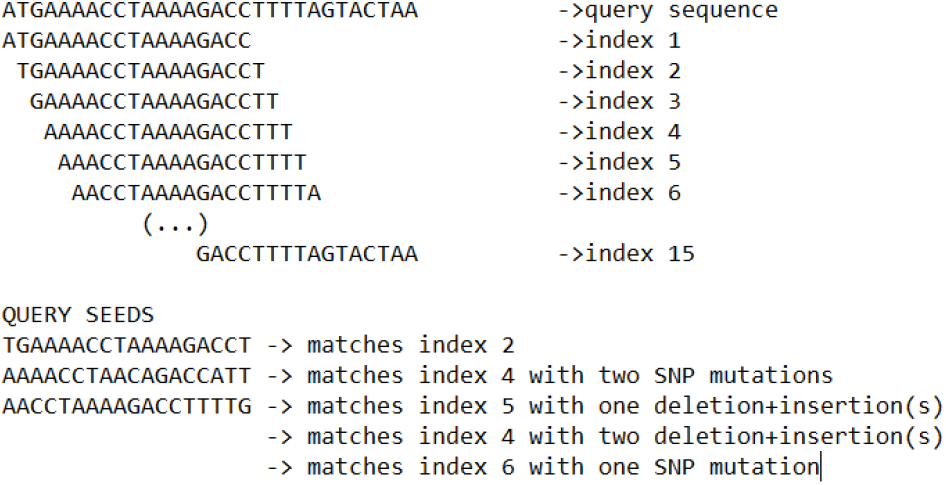
Matches for different query seeds.

So, we played with all these parameters and found a configuration that works well enough for our purposes and we let the user select the sensibility of the indexing tool to choose between some faster and less accurate modes or slower and more accurate ones.

### 3. Mapping matches in the database sequences

Once we have made a search for every seed, we map all these matches to the sequences in the database. Another parameter that can be configured is how many matches are returned per seed search. Usually, limiting that number increases performance and does not have any impact on the accuracy (for long enough sequences) given that even if a seed misses a match there are more seeds likely matching the same sequence.

### 4. Selecting regions with a higher density of matches

Finally, a window with the size of the query sequence is displaced along the sequences in the database and the window positions with more matches are selected. These are the regions for which the Smith-Waterman algorithm will be applied.

## III. BENCHMARKS

The algorithm is in general slower than BLAST in its actual version, but we are not sure it is because of intrinsic disadvantage and not because of not proper configuration and optimization of the code. But nonetheless, in most cases, it is not exponentially slower than BLAST.

**Fig. 4.**
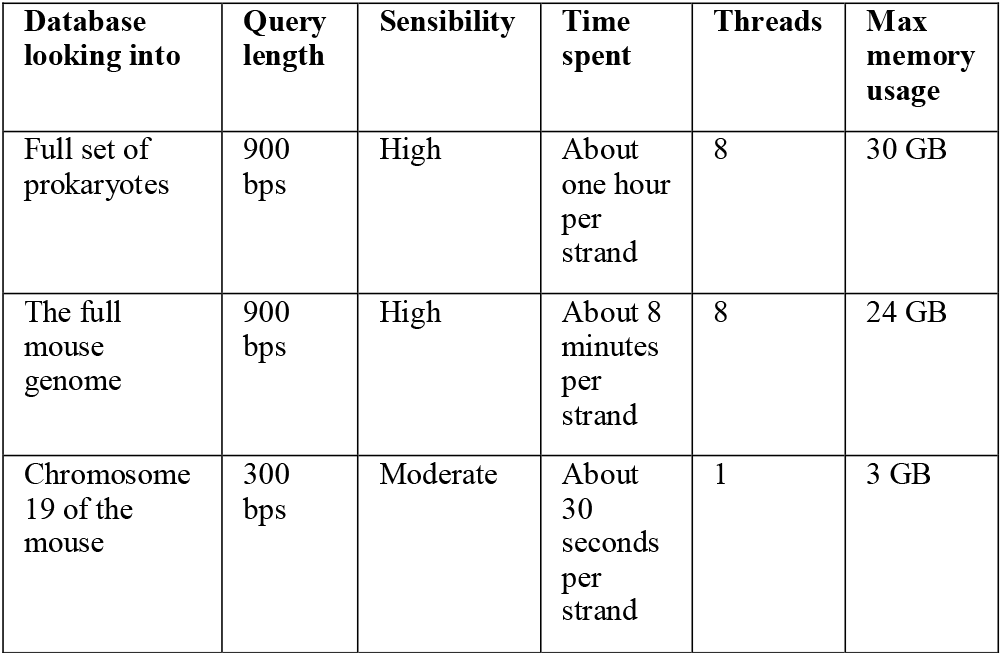
Time spent for making queries over different databases with a single computer (8 cores 30 GB of memory).

## IV. ADVANTAGES

We find that SLAST has advantages for some tasks.

### A. Finding homolog short non-conserved sequences

For short sequences not subject to conservation the accumulation of mutations can make related sequences hardly recognizable, especially for algorithms that focus on finding consecutive matches between base pairs, as BLAST does. Therefore, BLAST will return even shorter sequences as matches, these being so short that they could even be due to chance in some cases. BLAST does not compare the rest of the sequence.

In contrast, once SLAST finds a region with enough matching seeds, it tries to make an alignment of similar size to the query sequence. Therefore, even if the sequence is very eroded by mutations, the sequences with fewer mutations will still have less distance to the query than the most eroded ones. This way, you will get higher scores for closer matching results and lower scores for more eroded ones.

For example, if you make a search for AluSq10 sequences in the genome of the mouse, BLAST will return thousands of short matches on average 60 bps long, while the sequence used as query was 292 bps long.

**Fig. 5.**
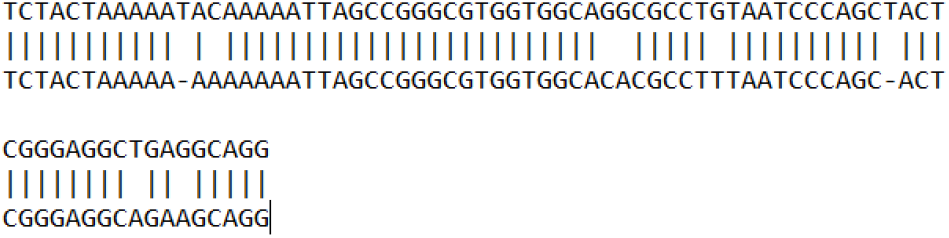
Pairwise alignment of fragment of AluSq10 query sequence and a BLAST match in chromosome 1 of the mouse

SLAST will return, in contrast, thousands of alignments like the one in fig. 6.

**Fig. 6.**
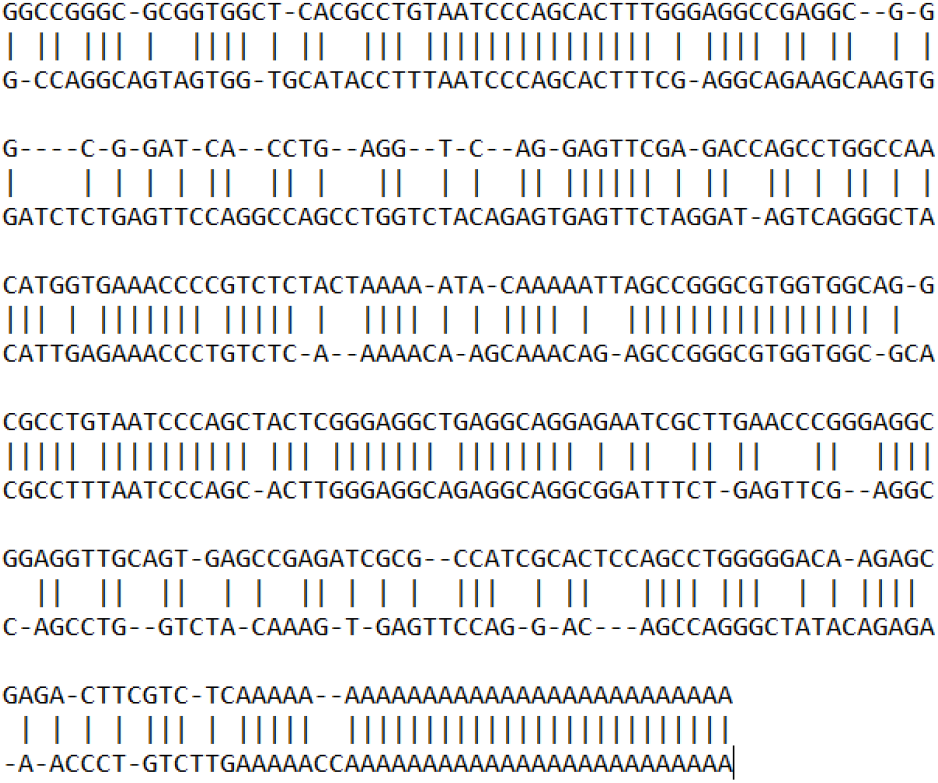
Pairwise alignment of AluSq10 query sequence and a SLAST match in chromosome 19 of the mouse

Note also how BLAST omits repetitive sequences like the final A’s which in some circumstances can enhance speed and limit useless matches but not in this case.

### B. Getting data for phylogenetic trees

Phylogenetic trees usually require a multiple alignment of sequences to compare. If these sequences have different sizes, the difference in size is filled with gaps. These gaps, therefore increase the distance between the sequences even if some bps in the original sequences were matching. That, therefore, takes to a less reliable tree.

Therefore, SLAST matching sequences are more useful for making phylogenetic trees than BLAST ones, especially when comparing a large number of short sequences and when manually curating the data is not a practical option.

## V. CONCLUSIONS

Despite we have not found evidence that a strategy consisting on applying the Smith-Waterman algorithm to regions with a high density of seed matches has, in most cases, any advantage over the usual seed and grow strategy, we find that in some cases, it does.

We need to investigate further for additional use cases for which our strategy could also have advantages.

On the other hand, we have not found evidence either that this strategy has less potential for designing this sort of tool (finding similar sequences in a database), being actually its only considerable drawback that it’s slower than BLAST. But we have not tested yet how the performance compares when applying FPGAs or GPUs to this strategy. That also is left for future research.

## Supporting information

SLAST results in mouse chromosome 19 for AluSq10 query

## VI. ACKNOWLEDGMENTS

The authors of this research adhere to the Agile Science Manifesto v 1.0 [3] and this document was written in concordance with its arguments.

## Notes

https://www.dnaservic.es/dnaresult?ob=2&min=0.5&id=5191&qid=BdZIQbHRnfIsdH0O&tkt=&code=

